# The soybean *Rhg1* amino acid transporter protein becomes abundant along the SCN penetration path and impacts ROS generation

**DOI:** 10.1101/2020.09.01.277814

**Authors:** Shaojie Han, John M. Smith, Yulin Du, Andrew F. Bent

## Abstract

*Rhg1* mediates soybean resistance to soybean cyst nematode. *Glyma.18G022400*, one of three resistance-conferring genes at the complex *Rhg1* locus, encodes the putative amino acid transporter AAT_Rhg1_ whose mode of action is largely unknown. We discovered that AAT_Rhg1_ protein abundance increases 7- to 15-fold throughout root cells penetrated by SCN. These root cells develop increased abundance of vesicles and larger vesicle-like bodies. AAT_Rhg1_ was often associated with these vesicles. AAT_Rhg1_ abundance remained low in syncytia (plant reprogrammed feeding cells), unlike the *Rhg1* α-SNAP protein whose abundance was previously shown to increase in syncytia. In *N. benthamiana*, if soybean AAT_Rhg1_ was present, oxidative stress promoted formation of larger macrovesicles and they contained AAT_Rhg1_. AAT_Rhg1_ was found to interact with GmRBOHC2, a soybean ortholog of Arabidopsis RBOHD previously found to exhibit upregulated expression upon SCN infection. Reactive oxygen species (ROS) generation was more elevated when AAT_Rhg1_ and GmRBOHC2 abundance were co-expressed. These findings suggest that AAT_Rhg1_ contributes to SCN resistance along the penetration path as SCN invades the plant, and does so at least in part by interactions with GmRBOHC2 that increase ROS production. The study also shows that *Rhg1* resistance functions via at least two spatially and temporally separate modes of action.

## INTRODUCTION

*Rhg1* of soybean (*Glycine max*) is a complex genetic locus that encodes novel mechanisms of disease resistance (Cook et al., 2012; Bayless et al., 2016; Mitchum, 2016). Soybean is one of the four largest world food crops and *Rhg1* is a central tool used for control of the most economically damaging disease of U.S. soybeans, the soybean cyst nematode (SCN, *Heterodera glycines*) (Niblack et al., 2006; Jones et al., 2013; Mitchum, 2016; Allen et al., 2017). The modes of action of *Rhg1* remain only partially understood.

Recently hatched J2 SCN migrate toward soybean root exudates and then penetrate soybean roots above the root cap in the zone of elongating cells (Endo, 1992). They then migrate intracellularly through root cortical cells, secreting bioactive effectors through their stylet to degrade cell walls and manipulate plant defense responses (Qin et al., 2004; Gheysen and Mitchum, 2011; Mitchum et al., 2013; Sato et al., 2019). The protrusible stylet is also is used directly to disrupt plant cell walls prior to cell penetration. When SCN reach a suitable site adjacent to the vascular cylinder they select an individual endodermis or endodermis-adjacent cortical cell upon which to feed. Over subsequent days the cell walls partially dissolve between an increasingly large cluster of dozens of adjacent root cells that also lose their large central vacuoles and become metabolically hyperactive, forming a multinucleate syncytium from multiple cytoplasmically merged cells (Jones, 1981; Fenoll et al., 1997; Kyndt et al., 2013). After 2-4 weeks of deriving nutrients from the host though a syncytium, the life cycle of a fertilized female SCN is completed by forming an egg-filled and durable cyst. Despite a number of previous studies (Mahalingam and Skorupska, 1996; Hermsmeier et al., 1998; Escobar et al., 2011; Kandoth et al., 2011), there is particularly incomplete knowledge regarding how plants respond during the early stages of infection as SCN penetrate and migrate through relatively new root tissues.

The structure of the complex *Rhg1* locus has been characterized across a wide array of soybean germplasm (Cook et al., 2012; Cook et al., 2014; Lee et al., 2015). Increased copy number of a four-gene block is a hallmark of resistance-conferring *Rhg1* haplotypes (Cook et al., 2012). “Peking”-type (*rhg1-a*) haplotypes typically carry three copies while “PI 88788”-type (*rhg1-b*) haplotypes often carry nine or ten copies of the ~30kb *Rhg1* four-gene block (Cook et al., 2012). *rhg1-a* generally must be combined with an appropriate allele of the unlinked *Rhg4* locus to achieve sufficient SCN resistance (Brucker et al., 2005; Liu et al., 2012; Mitchum, 2016). Until recent breeding efforts, most soybean accessions carried single-copy “Williams 82”-type *Rhg1_WT_* loci and were SCN-susceptible (Niblack et al., 2008; Lee et al., 2015; Bayless et al., 2019). In the U.S. market, over 95% of SCN-resistant soybean varieties now rely heavily on the PI88788-derived *rhg1-b* haplotype (Niblack et al., 2008; Tylka and Mullaney, 2015; Rincker et al., 2017)

Gene silencing and gene overexpression approaches have demonstrated contributions to SCN resistance by at least three of the four genes within the multicopy *Rhg1* segment (Cook et al., 2012; Liu et al., 2017; Butler et al., 2019). The resistance-contributing *Rhg1* genes include *Glyma.18G022400* (formerly named *Glyma18g02580*), which encodes a putative amino acid transporter hereafter referred to as *AAT*_Rhg1_, as well as *Glyma.18G022500* (formerly named *Glyma18g02590*), encoding a predicted α-SNAP (alpha-soluble NSF [N-ethylmaleimide–sensitive factor] attachment protein), and *Glyma.18G022700* (formerly named *Glyma18g02610*), encoding a protein with a WI12 wound-inducible protein domain (Cook et al., 2012). Several recent studies on *Rhg1*-encoded α-SNAP proteins have revealed their elevated abundance in syncytia and their cytotoxicity, which apparently poisons syncytium cells during the otherwise biotrophic plant-nematode interaction (Cook et al., 2012; Matsye et al., 2012; Cook et al., 2014; Bayless et al., 2016; Lakhssassi et al., 2017; Liu et al., 2017; Bayless et al., 2018; Bayless et al., 2019).

The products of the other two *Rhg1* genes that contribute to SCN resistance, *Glyma.18G022400* (AAT_Rhg1_) and *Glyma.18G022700* (WI12 protein), had been much less well characterized. There are no predicted amino acid polymorphisms in the products of those genes between SCN-susceptible *Rhg1_WT_* and the resistance-conferring low-copy *rhg1-a* and high-copy *rhg1-b* haplotypes. However, the higher copy numbers of those *Rhg1* genes result in constitutive elevation of their transcript abundance in SCN-resistant plants (Cook et al., 2014), and locus copy number has been shown to correlate with the strength of SCN resistance conferred by various *rhg1-a* and *rhg1-b* haplotypes (Cook et al., 2014; Lee et al., 2016; Yu et al., 2016; Patil et al., 2019).

The amino acids transported by AAT_Rhg1_, if any, are not known. The protein retains the sequence hallmarks of a bona fide amino acid transporter but has been recalcitrant in yeast and *Xenopus* oocyte experiments attempting to document transport of particular amino acids. A recent publication showed that overexpression of *Glyma.18G022400* (AAT_Rhg1_) in soybean can increase tolerance of toxic levels of exogenously supplied glutamate, and presented additional indirect evidence showing that AAT_Rhg1_ impacts glutamate abundance and transport (Guo et al., 2019). Endogenous jasmonic acid (JA) levels and JA pathway genes were also upregulated in AAT_Rhg1_ overexpression soybean lines (Guo et al., 2019). However, evidence is still lacking regarding mechanistic roles of AAT_Rhg1_ in SCN-soybean interactions. In gene expression studies, *Glyma.18G022400* transcripts were detected in syncytia cell samples captured by laser capture microdissection at 3, 6 and 9 days post-infection in both *rhg1-a* and *rhg1-b* (resistant) plants, but not in the susceptible lines. (Matsye et al., 2011).

Reactive oxygen species (ROS) are commonly produced during plant-pathogen interactions and act up- and downstream of various signaling pathways (Apel and Hirt, 2004; Camejo et al., 2016; Waszczak et al., 2018). Mechanisms that control ROS production during infections, and the impacts of ROS on infection outcomes, are finely nuanced and continue to be discovered. ROS generated during pattern-triggered immunity (PTI) act as important defense signal transduction molecules (Macho and Zipfel, 2014). ROS generated from extracellular and/or intracellular sources can accumulate to more toxic levels during the hypersensitive response, the programmed cell death response associated with effector-triggered immunity (ETI) (Zurbriggen et al., 2010). Plant ROS production can also contribute to disease susceptibility, for example in soybean interactions with necrotrophic *Sclerotinia sclerotiorum* (Ranjan et al., 2018). However, moderate levels of ROS can help limit the extent of cell death (Torres et al., 2005). This appears to be the case during colonization of Arabidopsis by beet cyst nematodes, for which ROS responses were shown to help limit plant cell death and enhance nematode growth (Siddique et al., 2014). In other cases, ROS have been shown to contribute to plant defense against plant parasitic nematodes (Simonetti et al., 2010; Kandoth et al., 2011; Kong et al., 2015; Pant et al., 2015; Teixeira et al., 2016; Habash et al., 2017; Labudda et al., 2018; Lee et al., 2018; Mei et al., 2018; Zhou et al., 2018; Yang et al., 2019b; Yang et al., 2019a; Chen et al., 2020; Hawamda et al., 2020; Labudda et al., 2020).

Plasma membrane-localized NADPH oxidases, encoded by respiratory burst oxidase homolog genes (*RBOH*), are key enzymes for pathogenesis-associated ROS generation (Averyanov, 2009). In soybean, the 17 *GmRBOH* genes were recently characterized by two separate groups (Ranjan et al., 2018; Liu et al., 2019). As part of that work, expression patterns of each *GmRBOH* gene in response to multiple biotic and abiotic stresses were analyzed using qRT-PCR and previously published microarray data (Liu et al., 2019), and one of the GmRBOHs was reported to be specifically induced by *Sclerotinia sclerotiorum* infection (Ranjan et al., 2018).

In Arabidopsis one particular RBOH, RBOHD, plays major roles in mediating ROS production during both PTI and ETI (Kadota et al., 2015). For example, AtRBOHD interacts with the FLS2 immune receptor complex, and is phosphorylated by BIK1 to enhance ROS generation that contributes to stomatal closure defense mechanisms against *Pseudomonas* bacteria (Li et al., 2014). PBL13 associates with and directly phosphorylates RBOHD and a newly discovered E3 ubiquitin ligase PIRE could interact with both RBOHD and PBL13 to stimulate degradation of RBOHD (Lee et al., 2020). RBOHD orthologs in multiple plant species have been reported to control PTI (Simon-Plas et al., 2002; Yoshioka et al., 2003; Trujillo et al., 2006; Kobayashi et al., 2007; Wong et al., 2007; Li et al., 2015). RBOHD contributes to defense against root knot nematodes (Teixeira et al., 2016), and reduced H_2_O_2_ production due to silencing of the tomato *RBOHD* homolog caused greater susceptibility (Zhou et al., 2018). In soybean, transcript abundance of the Arabidopsis *RBOHD* ortholog *Glyma.06G162300* (encoding GmRBOHC2) is significantly upregulated during SCN infection in both susceptible and resistant lines (Wan et al., 2015; Liu et al., 2019), but other aspects of GmRBOHC2 behavior during SCN infestations have not been characterized.

The present study examined soybean AAT_Rhg1_. We discovered unique accumulation of this protein along the SCN penetration path. Vesiculation occurred in root cortical cells penetrated during SCN migration and AAT_Rhg1_ was often present on those vesicles. Heterologous overexpression of AAT_Rhg1_ could induce vesiculation. Further, we found that AAT_Rhg1_ interacts with GmRBOHC2, a soybean ortholog of Arabidopsis RBOHD. Simultaneous overexpression of GmAAT_Rhg1_ and GmRBOHC2 elevates ROS production. The findings suggest that the *Rhg1*-encoded α-SNAP_Rhg1_ and AAT_Rhg1_ proteins contribute to SCN resistance through temporally, spatially and biochemically distinct mechanisms.

## RESULTS

### Soybean AAT_Rhg1_ protein abundance is elevated along the penetration path of soybean cyst nematode

The abundance of native AAT_Rhg1_ in soybean roots was qualitatively assessed via standard Western immunoblots using a custom antibody raised against a unique AAT_Rhg1_ peptide sequence (Figure 1). SCN-infested root regions or analogous regions from mock-inoculated controls were harvested at 4 days post-infection (dpi) from non-transgenic Wm82, Forrest, and Fayette cultivars, which respectively carry wild type *Rhg1* (single-copy/susceptibility-associated), or *rhg1-a* (3 copies of resistance-associated *Rhg1*), or *rhg1-b* (10 copies of resistance-associated *Rhg1*). Figure S1 confirms antibody recognition of the intended gene product. In non-infected roots, AAT_Rhg1_ protein abundance was low and similar in WT and low-copy *rhg1-a* samples but consistently greater in high-copy *rhg1-b* samples (Figure 1A). An obvious increase in AAT_Rhg1_ protein abundance was observed in SCN-infected samples of high-copy *rhg1-b* roots (Figure 1A). Any AAT_Rhg1_ abundance increases were more subtle for susceptible/wild-type and for low-copy *rhg1-a* roots. Figure 1B presents densitometric quantification of the immunoblot band intensities for four samples per treatment from two independent experiments.

**Figure 1.**
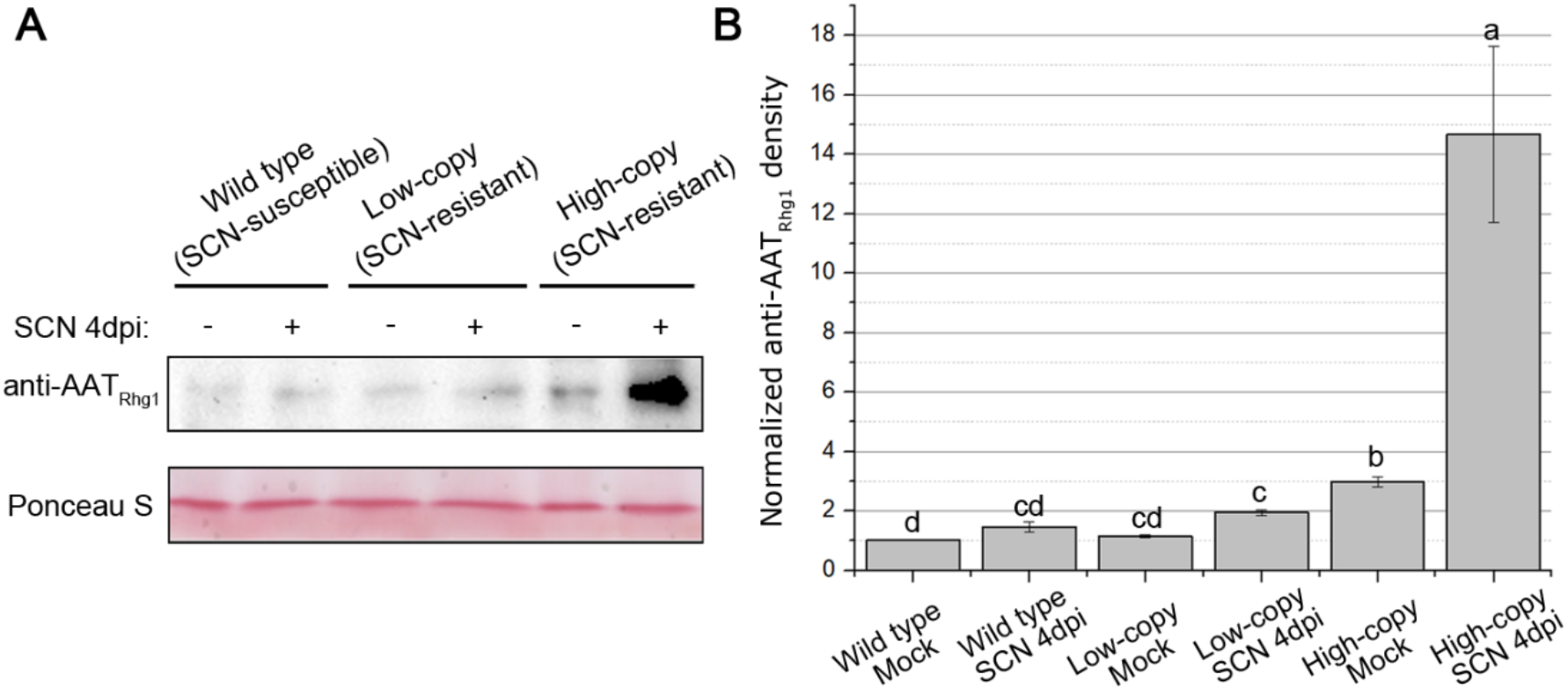
AAT_Rhg1_ protein abundance increase upon SCN infection in *rhg1-b* soybean roots. **(A)** Representative immunoblot showing the abundance of AAT_Rhg1_ protein in soybean lines that carry *Rhg1_WT_* (wild type), *rhg1-a* (low-copy), or *rhg1-b* (high-copy) haplotypes. Detached roots from the three varieties were either infected with an Hg Type 0 population of SCN (SCN +) or mock-inoculated (SCN -), and harvested at 4 dpi. Samples of SCN-infested root regions from four roots per treatment were pooled together for the immunoblot. Ponceau staining of the blotted membrane shown as check for equivalent loading of total protein. **(B)** Densitometry analysis of immunoblot AAT_Rhg1_ protein levels. n=4 for each treatment; data obtained from two biological replicates. Band intensity was normalized to the intensity for wild type mock treatment within each blot; mean ± SE are shown. Bars with the same letter are not significantly different (ANOVA Tukey analysis performed on non-normalized data, p < 0.05).

Transmission electron microscopy (TEM) immunogold detection experiments were conducted to provide cellular and subcellular-level resolution regarding AAT_Rhg1_ protein location and relative abundance. Previously, we used similar methods for *Rhg1* locus α-SNAP_Rhg1_ proteins and discovered more than ten-fold greater accumulation of α-SNAP_Rhg1_HC or α-SNAP_Rhg1_LC within syncytium cells (the root cells that comprise the differentiated SCN feeding site), relative to the surrounding cells (Bayless et al., 2016; Bayless et al., 2019). Using the anti-AAT_Rhg1_ antibody to detect native AAT_Rhg1_ in non-transgenic roots during SCN infection, we observed an entirely different pattern (Figure 2). Relative to adjacent cells, AAT_Rhg1_ protein abundance was elevated in penetrated root cells along the migration path of SCN. Figure 4 includes representative lower-magnification images showing the SCN body, SCN-penetrated root cells and adjacent normal root cells. Using higher magnification, the experiments of Figure 2 found 4.4-fold to 12.5-fold more immunogold labeled AAT_Rhg1_ in SCN-penetrated cells relative to a similar area in adjacent normal root cells, across three independent experiments. Increases were observed in both SCN-susceptible and two types of SCN-resistant cultivars (Figure. 2A, 2B). At 3 dpi, anti-AAT_Rhg1_ immunogold particle abundance values relative to adjacent normal cells were lowest for single-copy *Rhg1* (susceptible) roots (~4.4-fold elevation), moderate for low-copy *rhg1-a* (resistant) roots (~7.3-fold elevation), and the highest for high-copy *rhg1-b* (resistant) soybean roots (~11.1-fold elevation). However, at 7 dpi, all genotypes accumulated AAT_Rhg1_ to similar fold change levels (9.9, 11.0 and 12.5 fold-changes in single-copy, low-copy and high-copy resistant respectively). In all cases, the elevated abundance of AAT_Rhg1_ signal relative to nearby non-penetrated root cells within the same microscopy grid was statistically significant (Figure 2B).

**Figure 2.**
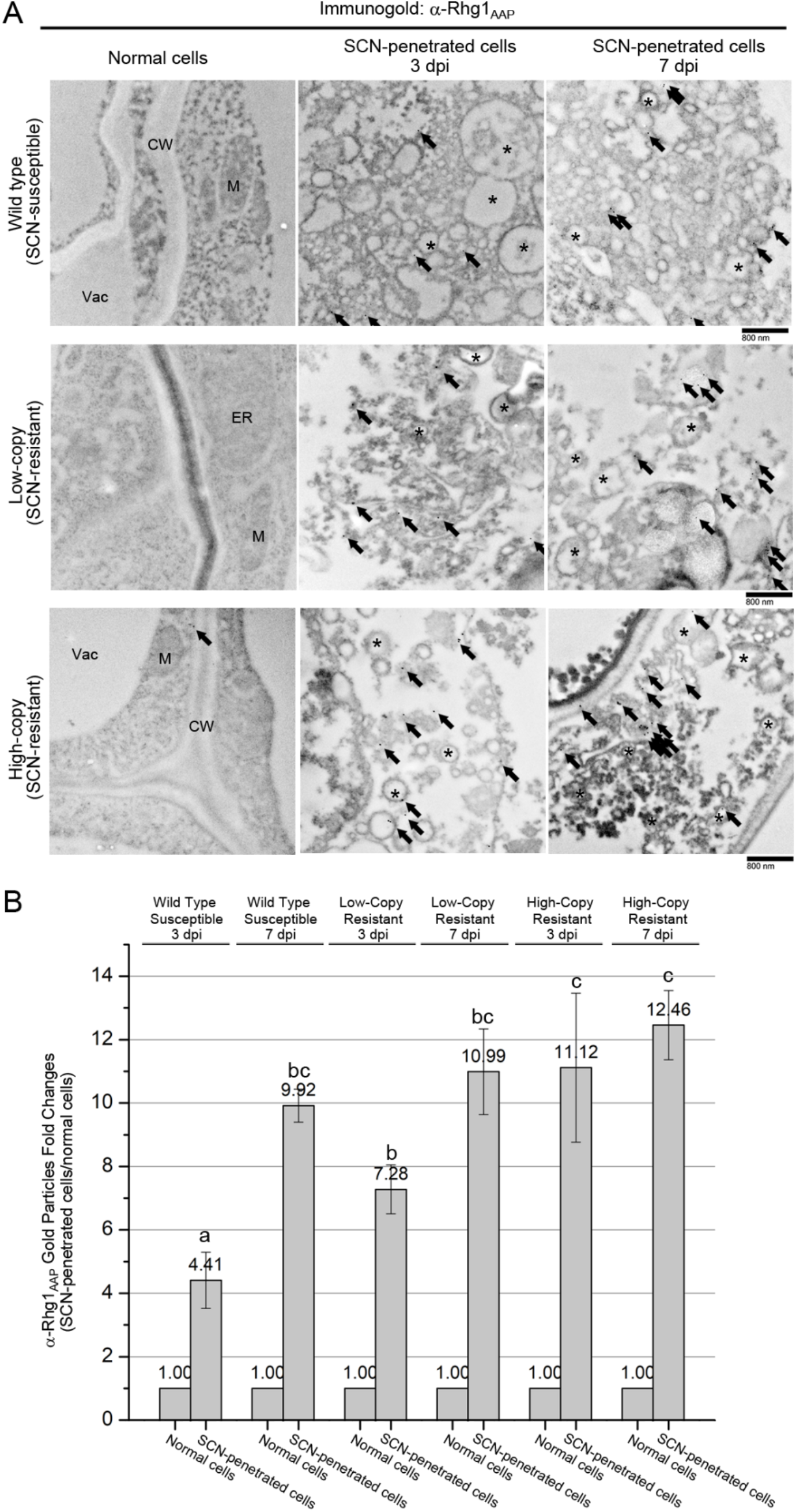
AAT_Rhg1_ abundance increases at SCN-penetrated root cells and is localized on vesicles and macrovesicles induced by SCN migration. **(A)** Representative electron micrographs taken at 15,000X magnification showing immunogold-labeled AAT_Rhg1_ (solid black particles) on vesicle membranes of SCN-penetrated cells in SCN infested roots of susceptible (upper panels), *rhg1-a* SCN resistant (low-copy, middle panels), and *rhg1-b* SCN resistant (high-copy, bottom panels) genotypes, at 3 dpi (middle column) and 7 dpi (right column), but not in normal cells (left column) from the same samples. Arrows indicate immunogold particles in each image. Asterisks highlight vesicle clusters in SCN-penetrated cells. CW, cell wall; M, mitochondrion; Vac, vacuole. Bars = 800 nm. **(B)** Number of AAT_Rhg1_ immunogold particles in SCN-penetrated cells relative to the highest number in a similar 2D area of an adjacent normal cell on the same grid (whose quantity is therefore 1.0 for each treatment). At least 30 images, from three independent experiments, were used to quantify AAT_Rhg1_ immunogold particle abundance for each treatment. Values are mean ± SE. Treatments marked with the same letter were not significantly different (ANOVA, P < 0.05).

**Figure 3.**
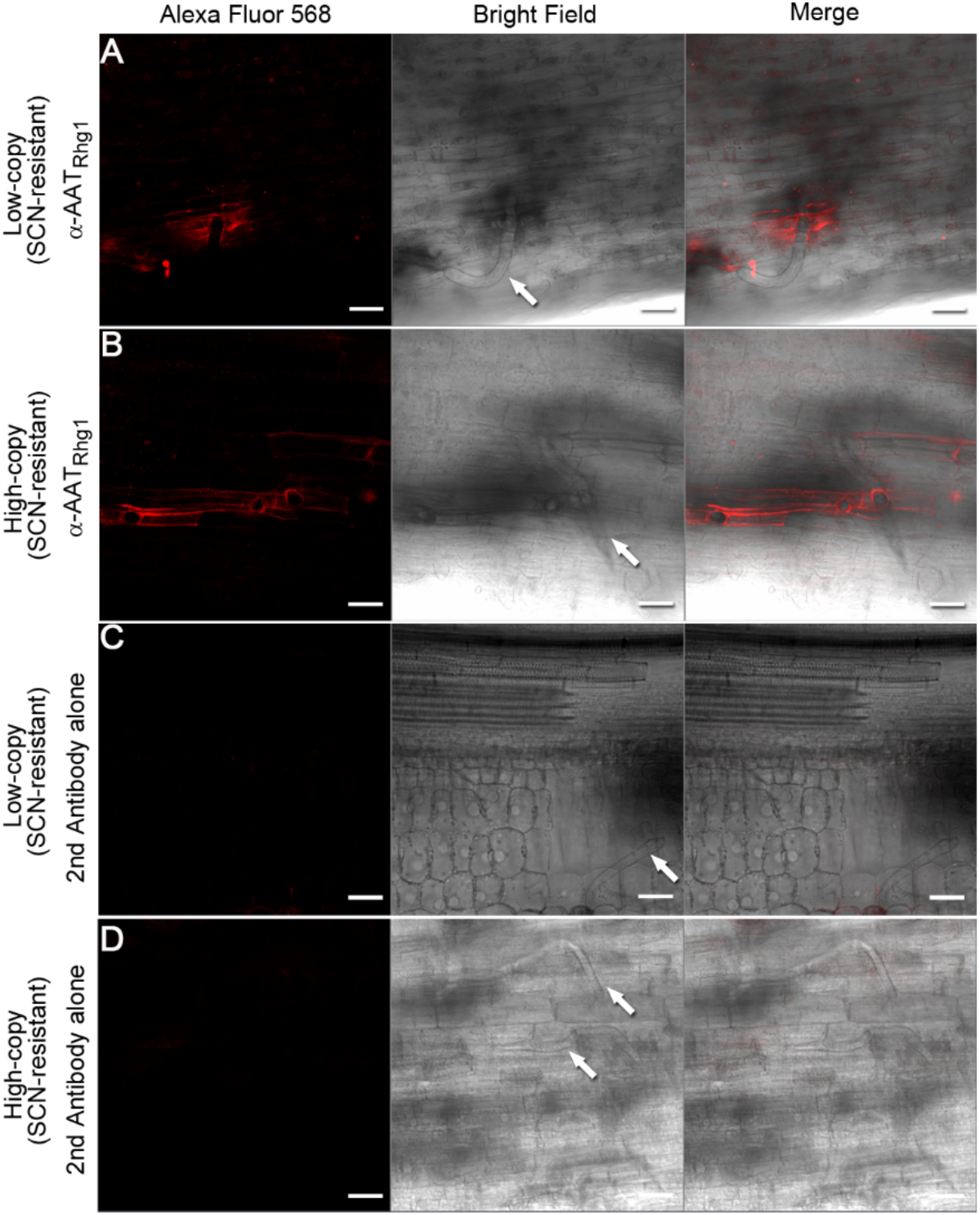
AAT_Rhg1_ abundance increase throughout cells penetrated by SCN. Representative confocal images of immunofluorescent AAT_Rhg1_ signal in SCN-infested roots of *rhg1-a* low-copy SCN resistant (A, C), and *rhg1-b* high-copy SCN resistant (B, D) soybean lines at 4 dpi. (A, B) anti-AAT_Rhg1_ antibody conjugated with secondary antibody, indicating AAT_Rhg1_ localization (red fluorescence, left column). (C, D) only secondary antibody was used, to serve as a control. T-PMT channel was used to collect the bright field/DIC images (middle column). Nematode bodies are indicated by white arrows. The right column contains merged images. Bars = 50 μm.

**Figure 4.**
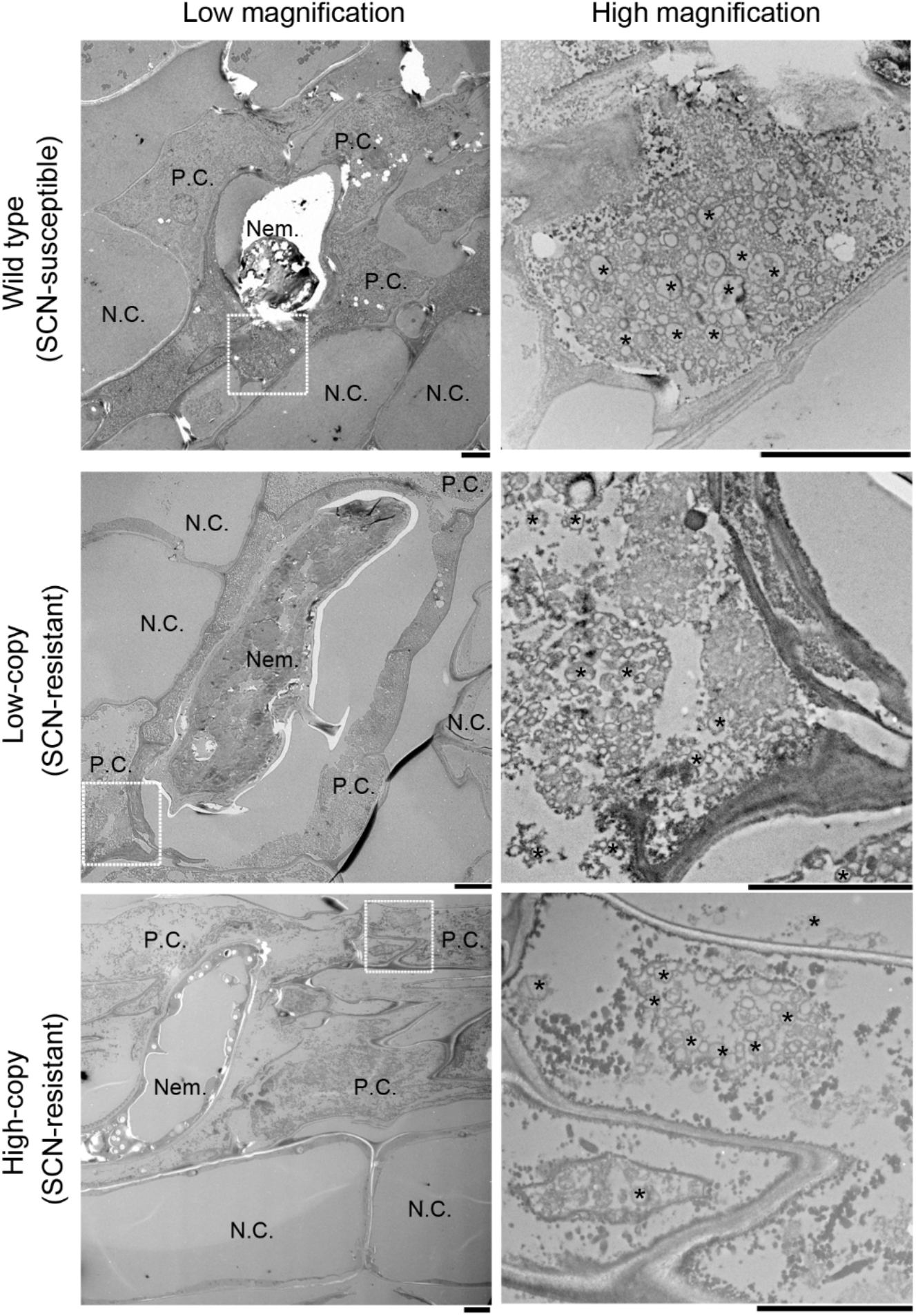
Substantial vesiculation in SCN-penetrated cells in SCN-infested roots of all tested soybean varieties. Electron micrographs showing root sections of SCN susceptible (top panels), *rhg1-a* low-copy SCN resistant (middle panels), and *rhg1-b* high-copy SCN resistant (bottom panels) haplotypes. Left column: SCN-penetrated cells (P.C.) surrounding a nematode body (Nem.); 710X magnification. Right column: further magnification (5600X images) of the corresponding area indicated by a white dot box on the left. Typical vesicle clusters are marked by an asterisk in those SCN-penetrated cells. Nem., Nematode; N.C., Normal cell; P.C., SCN-penetrated cell. Bars = 6 μm.

Anti-AAT_Rhg1_ immunogold particles were rarely found in mock-treatment samples of all the three genotypes tested (Figure S2A). Importantly, in both a susceptible and the two types of resistant varieties, anti-AAT_Rhg1_ immunogold particles were also rare in root syncytium cells (which are readily identifiable by the absence of a large vacuole, abundant presence of organelles and partially degraded cell walls) (Figure S2B). Syncytium cells are the primary site of accumulation of the *Rhg1*-encoded α-SNAP_Rhg1_ protein (Bayless et al., 2016).

In control experiments, no specific immunogold labeling could be found in vesicles or other compartments of SCN-penetrated cells when only the secondary antibody was used (Figure S2C). In further control experiments, competitive binding assays were conducted to confirm the antigen specificity of the anti-AAT_Rhg1_ antibody in EM/immunogold labeling use (Figure S3). The N-terminal 44 amino acid peptide that contains the antigen recognized by our custom AAT_Rhg1_ antibody was purified and pre-incubated with AAT_Rhg1_ antibody at a 1-fold or 10-fold molar excess before use on electron microscopy (EM) sections. Multiple adjacent tissue-sections from one identical region were examined on separate EM grids. The numbers of AAT_Rhg1_ immunolabel gold particles within the same penetrated cells were counted for sections probed with AAT_Rhg1_ antibody pre-treated with 1-fold or 10-fold molar excess antigen and compared with the particle numbers for tissue sections probed with AAT_Rhg1_ antibody not pretreated with peptide antigen. Results showed that both 1-fold and 10-fold molar excess of antigen binding significantly reduced the AAT_Rhg1_ immunogold signals within SCN-penetrated cells (Figure S3). This indicates specificity of the anti-AAT_Rhg1_ immunogold signal for the intended antigen in immunogold-labeled EM soybean root specimens.

We further confirmed this novel discovery of AAT_Rhg1_ abundance elevation in SCN-penetrated root cells using a separate method, confocal microscopy with immunofluorescent detection. This method allows broader visualization of AAT_Rhg1_ distribution within root samples. Roots of non-transgenic SCN-resistant soybean varieties Forrest (*rhg1-a*) and Fayette (*rhg1-b*) were inoculated with 200 J2 SCN per root. Four days after inoculation, SCN-infected root regions were chemically fixed. The *in situ* location and abundance of native AAT_Rhg1_ protein were monitored by secondary detection of the anti-AAT_Rhg1_ antibody using an Alexa Fluor 568 dye-conjugated anti-rabbit IgG antibody. Under bright-field illumination, the SCN-penetrated cells could be readily identified due to the visible nematode body or the round hole caused by SCN-penetration (Figure 3). Confocal fluorescence microscopy detected anti-AAT_Rhg1_ antibody only in cells that had been penetrated by a nematode (Figure 3). Signal was detected throughout the entire cell rather than only at the site of penetration. Similar results were obtained for roots of Forrest (*rhg1-a*) and Fayette (*rhg1-b*) plants (Figure 3). No specific signal was detected when secondary fluorescent antibody alone was used (Figure 3).

Taken together, the above results indicate that SCN-penetrated cells undergo a substantial increase in the abundance of the *Rhg1*-encoded amino acid transporter-like protein (AAT_Rhg1_, *Glyma.18G022400* gene product). At the early 3 dpi infection stage the level of AAT_Rhg1_ accumulation within SCN-penetrated cells positively correlated with *Rhg1* copy number, with the ten-copy *rhg1-b* soybean variety accumulating the most AAT_Rhg1_.

### AAT_Rhg1_ abundance increase not observed upon wounding with needle

To test whether AAT_Rhg1_ is a general wound-inducible protein, roots of Fayette (*rhg1-b*) were penetrated with 100 μm diameter microneedle. After three days, wounded root regions were isolated and chemically fixed. Then as in the previous section, TEM immunogold labelling experiments, and separate confocal microscopy with immunofluorescent detection experiments, were performed using anti-AAT_Rhg1_ antibody. The mechanical damage caused by the microneedle did not elicit signal accumulation in or around the site of needle penetration (Figure S4). Although these experiments do not exclude the possibility that the particular forms or patterns of physical damage caused by the nematode can elicit elevated abundance of AAT_Rhg1_ signal, the experiments do provide evidence that simple physical penetration of the root cortex is not on its own sufficient to induce elevated abundance of AAT_Rhg1_.

### Vesicle abundance is elevated along the penetration path of soybean cyst nematode and AAT_Rhg1_ protein accumulates on those vesicles

Independent of immunogold label detection, the above-described TEM images revealed a second observation: a strong increase in the abundance of subcellular vesicles in those cells that had been penetrated by SCN (Figure 4; see also Figure 2). This increase in vesicle abundance was evident in susceptible roots as well as low-copy *rhg1-a* and high-copy *rhg1-b* soybean roots (Figure 4). In addition to numerous vesicles in the ~50-500 nm size range (similar to or larger than common transport or secretory vesicles), some 1-2 μm diameter “macrovesicles” were also present. Compared to the adjacent non-penetrated root cortical cells, which retain their large central vacuole, the cells directly surrounding the nematode body showed distinct morphology changes. First, the large central vacuoles were shrunken or otherwise replaced by a nematode body. Second, there were numerous vesicles clustered within the remaining cytoplasm. Third, organelles like mitochondria, ER or Golgi were rarely observed (Figure 4). In all types of soybean roots tested, close inspection of immunogold labeling showed accumulation and co-localization of the AAT_Rhg1_ protein onto those vesicles formed in penetration cells but not in adjacent normal cells (see for example Figure 2). The results indicate that SCN-penetrated cells undergo a substantial accumulation of vesicles, and that AAT_Rhg1_ protein accumulates on those vesicles.

### Overexpression of AAT_Rhg1_ in *N. benthamiana* induces vesiculation and AAT_Rhg1_ protein accumulates on those vesicles

As one means of investigating impacts of AAT_Rhg1_ protein accumulation in planta, we overexpressed soybean AAT_Rhg1_ in *N. benthamiana* leaves. GFP-tagged AAT_Rhg1_ or a GFP-only control, driven by a double CaMV 35S promoter, were transiently expressed in *N. benthamiana* leaves by agroinfiltration. 72 hrs after agroinfiltration, the localization of GFP-AAT_Rhg1_ was analyzed by confocal microscopy. The fluorescence signal for both GFP-AAT_Rhg1_ and GFP was readily detectable over background. Interestingly, in addition to its distribution throughout the plasma membrane, GFP-AAT_Rhg1_ was present in the form of multiple primarily cytoplasmic puncta (small spots) as well as large hollow (peripherally fluorescent) or solidly green-fluorescent vesicle-like structures (Figure 5). These large hollow vesicles were of various sizes from ~0.58 μm to 5.5 μm diameter, and clearly did not overlap with chloroplasts (Figure 5A). As expected, the GFP control mainly localized diffusely throughout the cytoplasm.

**Figure 5.**
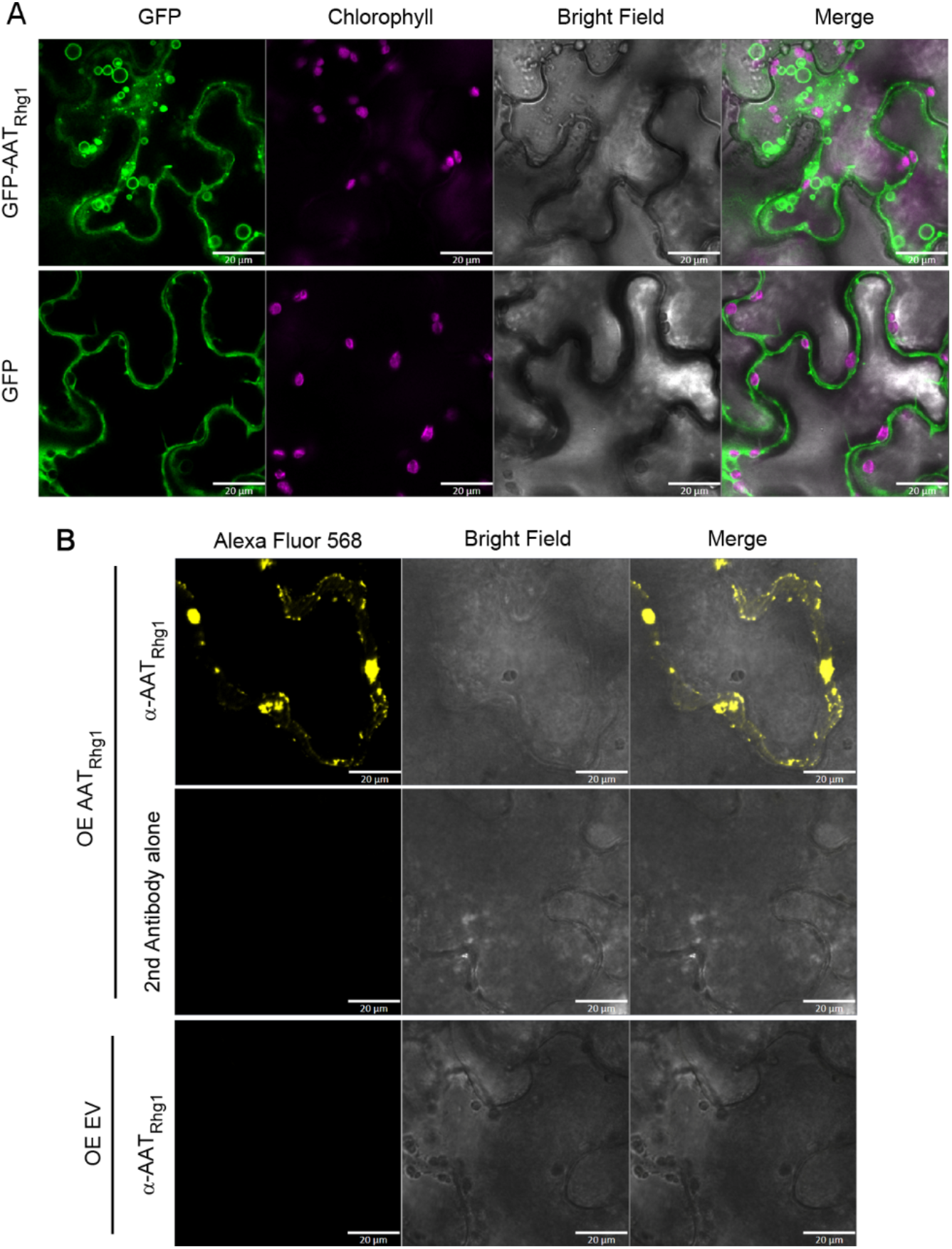
Soybean AAT_Rhg1_ localizes to specialized macrovesicles in *N. benthamiana*. **(A)** Top row: GFP-tagged GmAAT_Rhg1_ transiently expressed from a CaMV 35S promoter by agroinfiltration into *N. benthamiana* leaves. Left column: GFP-AAT Rhg1 localized on vesicles of sizes ranging from less than 1 μm (arrowhead) to ~6 μm (arrow). Chlorophyll signal (second column from left) and bright field image (third column from left) are from the same imaging layer. Merged images (right column) show that the AATRhg1-containing vesicles were independent of chloroplasts. Lower row: GFP alone expressed similarly as a control. Experiments were replicated on three separate dates with similar results. Bar = 20 μm. **(B)** Immunofluorescent stain confocal imaging showing that untagged GmAAT_Rhg1_ also accumulates in puncta of various sizes. *N. benthamiana* leaves expressing untagged GmAAT_Rhg1_ were immunostained with anti-AAT_Rhg1_ antibody then probed with secondary antibody conjugated to Alexa Fluor 568 (Top row). Leaf samples expressing the same GmAAT_Rhg1_ construct immunostained with secondary antibody alone (middle row) or samples expressing empty vector immunostained with both anti-AAT_Rhg1_ antibody and the secondary antibody (bottom row) served as controls. Images were acquired under the same settings across all three rows. Untagged soybean AAT_Rhg1_ formed puncta with sizes ranging from less than 1 μm (arrowhead) to ~6 μm (arrow) (top panels). Bar = 20 μm.

Because the localization of fluorescent protein-tagged proteins does not always reflect the authentic location of the untagged protein, we conducted additional immunolocalization analyses using untagged AAT_Rhg1_. *N. benthamiana* leaves transiently expressing soybean AAT_Rhg1_ or empty vector were chemically fixed at 72 hpi. Samples were incubated with the anti-AAT_Rhg1_ antibody followed by secondary incubation with Alexa Fluor 568 dye for immunofluorescence confocal microscopy. Other samples expressing AAT_Rhg1_ were incubated with secondary Alex Fluor 568 dye alone as a control. AAT_Rhg1_ immunofluorescent signal was detected in cells expressing AAT_Rhg1_ and not in empty vector controls (Figure 5B). The signal was specific to the primary anti-AAT_Rhg1_ antibody as the secondary antibody alone control did not show any signal (Figure 5B). Interestingly, the specific immunofluorescent signal again accumulated in puncta spots and in vesicle-like structures with an apparent size ranging from 0.94 μm to 6.6 μm. Unlike the localization of GFP-AAT_Rhg1_, AAT_Rhg1_ immunolocalization also showed rare cytoplasmic localization.

To associate the observed GFP-AAT_Rhg1_-containing macrovesicles with defined cellular structures, we used confocal laser scanning microscopy to test for co-localization of GFP-AAT_Rhg1_ and four organelle markers in *N. benthamiana* leaves (Nelson et al., 2007) (Figure S5). GFP-AAT_Rhg1_ was coexpressed with RFP-tagged markers for Golgi (Golgi-RK), ER (ER-RK), plasma membrane (PM-RK) or plastids (plastids-RK). Upon coexpression with each of the organelle markers, GFP-AAT_Rhg1_ displayed the same macrovesicle localization that GFP-AAT_Rhg1_ alone showed in Figure 5. Partial co-localization was observed for GFP-AAT_Rhg1_ and the ER marker, and for GFP-AAT_Rhg1_ and the plasma membrane marker (Figure S5). GFP-AAT_Rhg1_ did not show co-localization with the Golgi or plastid markers.

Together, the immunofluorescence and green fluorescent-protein tagging showed that overexpressing AAT_Rhg1_ causes plant cells to accumulate vesicles and macrovesicles, many of which carry AAT_Rhg1_.

### H_2_O_2_ accumulation in cells around the SCN at early infection stage

ROS signaling can be a mediator of plant defenses against pathogens and ROS production in roots during cyst nematode infections has been studied using multiple approaches. We examined the spatial pattern of ROS accumulation as SCN migrate through roots. ROS production was monitored in susceptible and in *rhg1-a* and *rhg1-b* SCN-resistant varieties, using the hydrogen peroxide probe 2’, 7’-dichlorodihydrofluorescein diacetate (H_2_DCFDA). Roots were examined at 3 days post-infection, a time at which nematode migration is still occurring but some nematodes have initiated feeding and other infections have terminated. We found that H_2_O_2_ was induced by SCN in root cells of all the three types tested, but not in the mock treatments (Figure 6). Interestingly, only a portion of root cells around the SCN (indicated by arrows in Figure 6A) had strong H_2_DCFDA fluorescent signals. Scattered lesions (dead root cells) in the area of nematode invasion are commonly observed a few days after initial root exposure to SCN, but neither the lesions (arrowhead in Figure 6A) nor the SCN (arrow with stem) showed elevated ROS signals. Reproducibly, there were more cells with positive H_2_DCFDA fluorescent signals in *rhg1-b* high-copy variety than the *rhg1-a* low-copy variety or the wild type. For each fluorescent image, ImageJ was used to divide the area of H_2_DCFDA fluorescent signals by the total root area (with the SCN body and the space outside the root tissue eliminated) to calculate the percent area with ROS (Figure 6B). Compared to the wild type mock treatment, the relative ROS accumulation was the lowest for wild type single-copy *Rhg1* (susceptible) roots (~6.8-fold elevation), moderate for low-copy *rhg1-a* (resistant) roots (~15.8-fold elevation), and the highest for high-copy *rhg1-b* (resistant) soybean roots (~34.6-fold elevation). There were no significant differences between all the mock treatments of the three varieties (Figure 6B).

**Figure 7.**
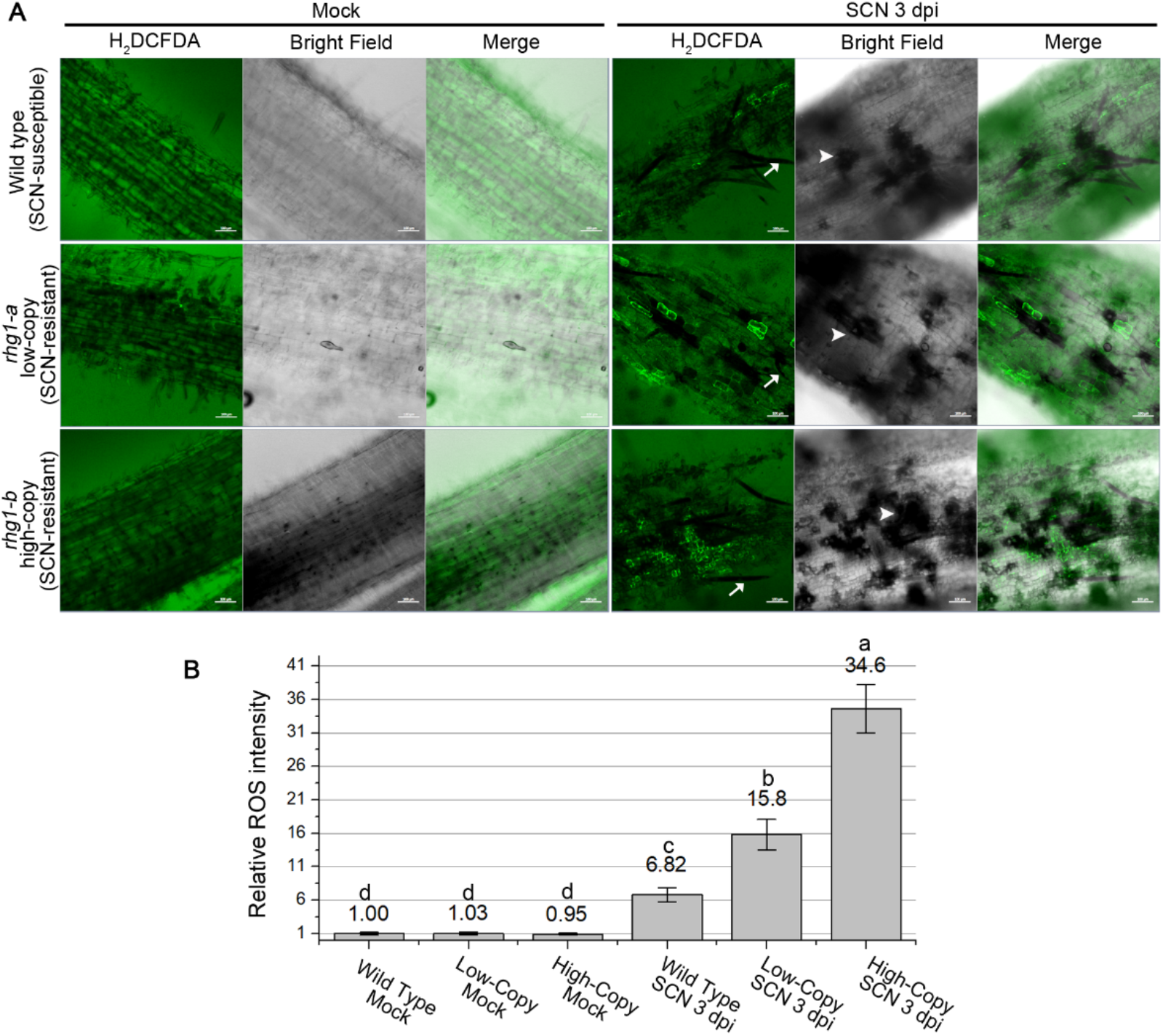
More cells accumulate ROS in the SCN infection zone of resistant roots. **(A)** Representative 3 dpi confocal images of mock-treated (left) and SCN-infected (right) soybean roots carrying wild type (top row), *rhg1-a* (middle row), and *rhg1-b* (bottom row) haplotypes, after incubation in H2DCFDA. H2DCFDA channel (green color) reports endogenous ROS production; bright field images shown in grayscale. Arrows with stem point to representative SCN body; arrowheads point to representative lesions caused by SCN infection. Bar = 100 μm. **(B)** Quantification of ROS-producing cells in soybean roots with or without SCN infection as in (A). Relative extent of ROS production was calculated by total H2DCFDA fluorescent area divided by total root cell area with fluorescent background and then normalized to wild type mock control treatment. For each treatment, at least 16 images of 8 independent roots from two independent experiments were used for calculation. Error bars indicate standard error of the mean. Treatments with the same letter are not significantly different (ANOVA, P < 0.05).

### ROS accumulation can enhance endocytosis-associated accumulation of AAT_Rhg1_-containing vesicles

Because soybean roots present a challenging experimental system for transient gene expression and for confocal imaging, *N. benthamiana* leaves were used to initiate investigation of the potential interaction of AAT_Rhg1_ with ROS. *N. benthamiana* leaves transiently expressing GFP-AAT_Rhg1_ or GFP alone as a control were infiltrated with 20 μm methyl viologen (MV; paraquat). MV is an inhibitor of photosynthetic electron transport chains that induces elevation of ROS in plant cells (Han et al., 2015). Eight hours after MV treatment, the leaf apoplast was infiltrated with FM4-64 and then imaged 30 mins later. Confocal live imaging of FM4-64 dye uptake is a standard technique to monitor vesicle dynamics in endocytic pathways (Bolte et al., 2004). As noted above, expression of GFP-AAT_Rhg1_ (in the absence of MV) led to the accumulation of green fluorescent puncta as well as green fluorescent vesicle-like structures (Figure 7, GFP column of images). Expression of GFP-AAT_Rhg1_ in the presence of MV led to the accumulation of more and larger green fluorescent vesicle-like structures with diameters of approximately 6-10 μm (arrow heads, Figure 7). Within some large vesicles multiple small round vesicles with GFP-AAT_Rhg1_ fluorescent signals were present, suggesting that the larger vesicles had formed through fusion of multiple small vesicles.

**Figure 7.**
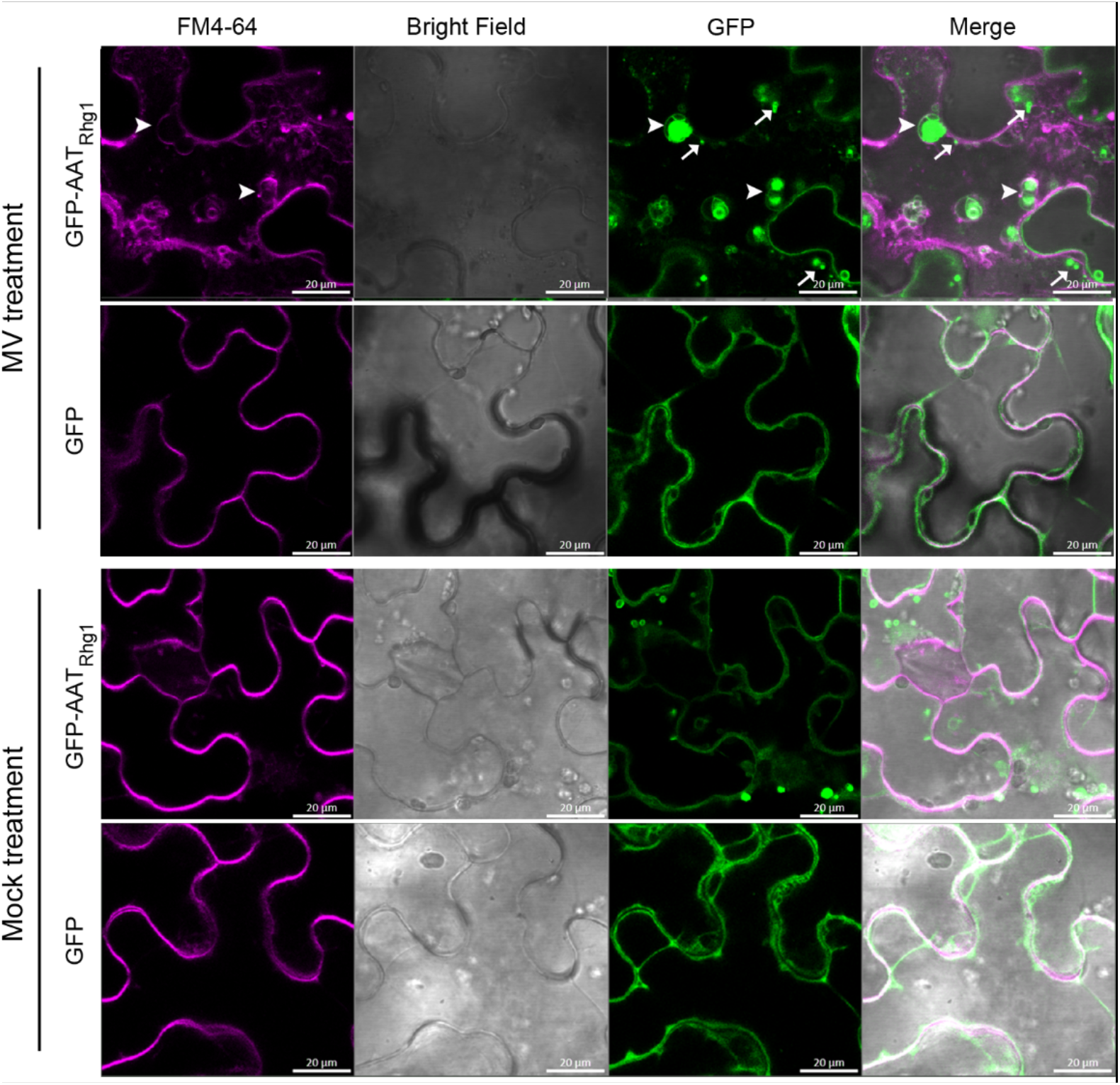
ROS stress leads to endocytosis-associated fusion of AAT_Rhg1_-containing vesicles. Representative confocal images showing, upon cellular ROS stress, AATRhg1-containing vesicles fused into larger vesicles through an endocytosis pathway in *N. benthamiana* leaf cells expressing GFP-AAT_Rhg1_ (top row) but not in cells expressing GFP control (second row). MV treatment: methyl viologen (inducer of superoxide and other ROS). Transiently transformed *N. benthamiana* leaves were treated with 20 μm MV or mock treatment for 8 hrs, then stained with the endocytic tracer FM4-64 30 min prior to confocal imaging. Arrows with stem point to representative puncta; larger arrowheads point to representative larger macrovesicle-like agglomerations of GFP-AATRhg1. Areas of overlap between the magenta and green signal are lighter (more white) in the merged image (right column). Bar = 20 μm.

Merging of FM4-64 and GFP images showed that FM4-64 stained membranes co-localized with the fused large vesicles (indicated by GFP fluorescence, arrow heads, Figure 7) but not with smaller GFP-AAT_Rhg1_-containing vesicles (arrows, Figure 7). This suggests that endocytic vesicles fuse with and possibly promote the agglomeration of GFP-AAT_Rhg1_ into larger clusters. The small GFP-AAT_Rhg1_ vesicles may have formed prior to the addition of FM4-64, or might not be derived from endocytic events. To reiterate, we found in *N. benthamiana* that upon MV-induced oxidative stress, smaller AAT_Rhg1_-containing vesicles more often fuse into larger macrovesicles. These fused larger vesicles could be stained by a 30-min FM4-64 treatment, indicating that this fusion is associated with an endocytic internalization process. This endocytic vesiculation response to ROS stimuli, which was observed in the presence of elevated AAT_Rhg1_ expression (Figure 7), was reminiscent of our electron micrograph discoveries that SCN-penetrated cells in soybean roots form larger AAT_Rhg1_-containing vesicles that can contain multiple small AAT_Rhg1_-containing vesicles (Figure 2).

### AAT_Rhg1_ interacts on vesicles with GmRBOHC2, an SCN-responsive NADPH oxidase homolog

We hypothesized that AAT_Rhg1_ physically interacts with one or more previously discovered defense-associated proteins, and used *N. benthamiana* agroinfiltration and co-immunoprecipitation (co-IP) to conduct in planta tests for interactions. A small number of potential AAT_Rhg1_ interactors were selected from the published literature regarding SCN-elicited responses as well as some SCN effectors. GmRBOHC2 was identified and then found upon testing to be an AAT_Rhg1_ interactor (Figure 8). *Glyma.06G162300*, which encodes GmRBOHC2, is a soybean ortholog of the gene encoding Arabidopsis RBOHD, an extensively studied mediator of the ROS burst associated with pathogen infection and wounding in Arabidopsis (Miller et al., 2009; Li et al., 2014; Lee et al., 2020). In genome-wide expression profiling, *Glyma.06G162300* transcript abundance was significantly upregulated after SCN infection in both susceptible and resistant lines, with greater abundance in resistant lines at 3 dpi (Wan et al., 2015; Liu et al., 2019). We co-expressed epitope-tagged GmRBOHC2-MYC with GFP-tagged AAT_Rhg1_ (or GFP-only control) in *N. benthamiana* leaves. Plant extracts were taken 60 hrs. After agroinfiltration, immunoprecipitated with anti-GFP antibody, the products were separated by SDS-PAGE, and protein blots were then probed with anti-MYC antibody. GmRBOHC2 co-immunoprecipitated with GFP-AAT_Rhg1_ but not with GFP alone (Figure 8A).

**Figure 8.**
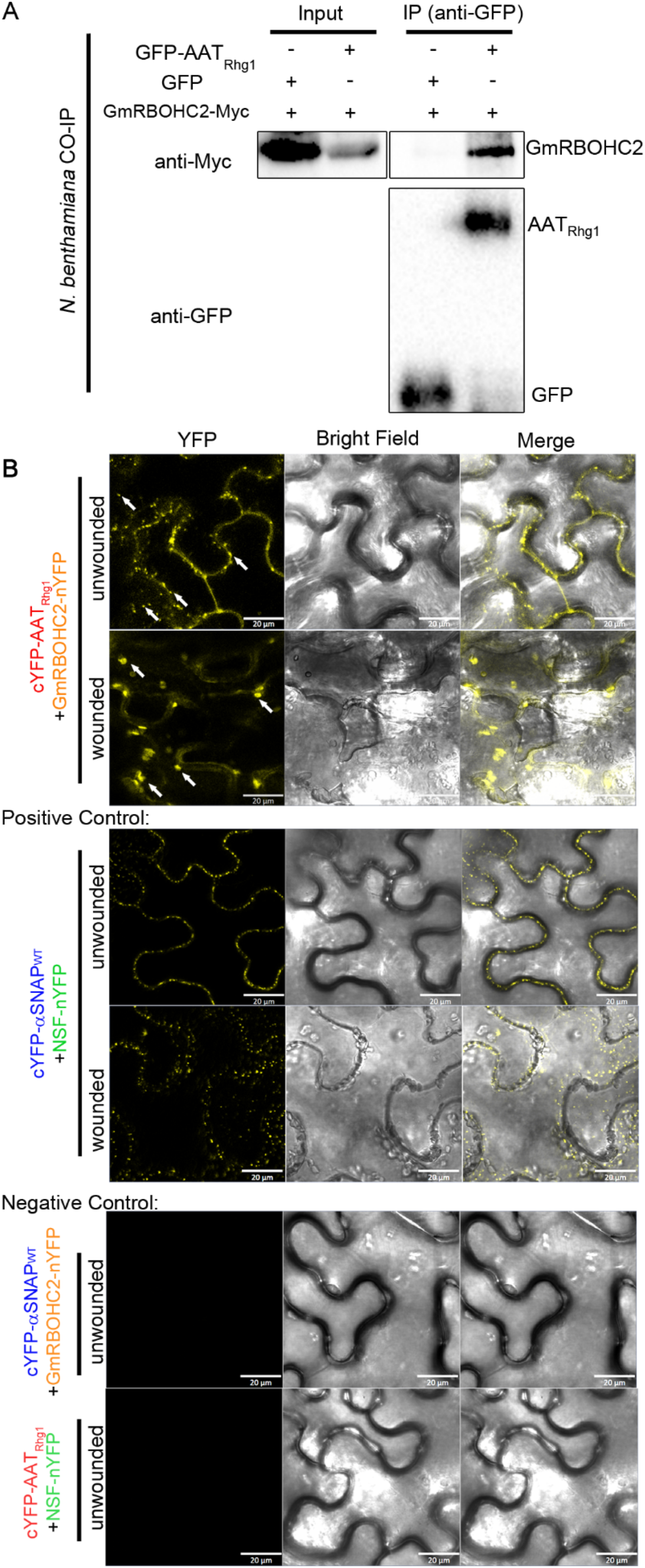
AAT_Rhg1_ and GmRBOHC2 interact in plants; wound treatment changes their interaction sites from small to large vesicles. **(A)** Immunoblots showing co-immunoprecipitation of AAT_Rhg1_ with soybean RBOHC2. Myc tagged GmRBOHC2 was co-expressed with GFP-AAT_Rhg1_ in *N. benthamiana* leaves by agroinfiltration. GmRBOHC2 co-expressed with GFP alone served as a negative control. IP (anti-GFP): leaf lysates harvested at 72 hours post infiltration were immunoprecipitated with anti-GFP beads and immunoprecipitates were assessed by immunoblotting using anti-Myc (right column, top panel) and anti-GFP antibodies (right column, bottom panel). Input: presence of GmRBOHC2-Myc in total protein in each treatment was confirmed (left blot). **(B)** Representative images from BiFC assays showing the localization of AAT_Rhg1_ interaction with GmRBOHC2 in plants, with or without wounding treatment. GmRBOHC2-nYFP was co-expressed transiently with cYFP-AAT_Rhg1_ in *N. benthamiana* leaves under unwounded conditions or wounded conditions (top two panels). As a positive control, GmNSF-nYFP and cYFP-α-SNAP_Rhg1_WT were coexpressed similarly (middle two panels). As a negative control the same constructs were coexpressed with different pairing (bottom two panels). YFP fluorescence indicated by yellow color (left column) was detected from epidermal cells. In cells coexpressing cYFP-AAT_Rhg1_ and GmRBOHC2-nYFP, complemented fluorescence signal was detected in small vesicles (indicated by white arrows in first panel) in the unwounded condition, or in large vesicles (white arrows in second panel) in wounded cells. The experiments were repeated on three separate dates with similar results. Bars =20 μm.

To validate these results and to investigate the cellular location of GmRBOHC2-AAT_Rhg1_ interaction, bimolecular fluorescence complementation (BiFC) experiments were carried out. Stringent positive and negative controls are necessary for BiFC experiments (Kudla and Bock, 2016); we used the known protein interaction partners NSF and α-SNAP_Rhg1_WT of soybean (Bayless et al., 2016) for this purpose. NSF and α-SNAP interact in vitro and in vivo, where they participate in the disassembly of SNARE protein bundles that are associated with vesicle trafficking, including at the plasma membrane (Zhao et al., 2015). Here, cYFP-tagged AAT_Rhg1_ was transiently coexpressed in *N. benthamiana* leaves with either RBOHC2-nYFP or the negative control NSF-nYFP. cYFP-tagged α-SNAP_Rhg1_WT was coexpressed with RBOHC2-nYFP to serve as another negative control. cYFP-α-SNAP_Rhg1_WT and NSF-nYFP were co-transformed within the same leaf as the above samples to serve as a positive expression control for the negative controls. We observed positive interaction signals, indicated by yellow fluorescence, only upon coexpression of cYFP-AAT_Rhg1_ and GmRBOHC2-nYFP, as well as for the positive control cYFP-α-SNAP_Rhg1_WT + NSF-nYFP (Figure 8B). Interestingly, the interaction signals were localized on small vesicle-like puncta within the cytoplasm, which was consistent previous AAT_Rhg1_ localization results (Figure 5). The known vesicle trafficking contributors α-SNAP and NSF also interacted in vesicle-like puncta in our BiFC assay (Figure 8B). The other control combinations did not give yellow fluorescence under the same confocal detection settings (Figure 8B).

As shown above (Figures 5 and 6), AAT_Rhg1_ localized onto larger fused vesicles under MV-induced ROS stress when overexpressed in *N. benthamiana* leaves. To test whether the co-localization pattern of AAT_Rhg1_GmRBOHC2 changed under similar stress, a hemostat wounding method was used after 60 hr co-expression of cYFP-AAT_Rhg1_/GmRbohC2-nYFP or cYFP-α-SNAP/NSF-nYFP control. After wounding, the leaves were left for 30 mins in the air before confocal microscopy. After this treatment, the reconstituted YFP signal indicating interaction of cYFP-AAT_Rhg1_ and GmRbohC2-nYFP was shifted toward larger vesicles (Figure 8B). The α-SNAP/NSF interaction signal remained on similarly sized vesicles with or without wounding treatment (Figure 8B). Hence the ROS-generating GmRBOHC2 protein, previously shown to be transcriptionally upregulated during SCN infection, interacted with AAT_Rhg1_ in small vesicles in normal conditions and in larger vesicles after wounding.

### Simultaneous elevation of AAT_Rhg1_ and GmRBOHC2 abundance causes ROS production

Having discovered that AAT_Rhg1_ and GmRBOHC2 physically interact in planta, experiments were then carried out to determine if AAT_Rhg1_ alters ROS generation in concert with GmRBOHC2 (Figure 9). GmRBOHC2 and AAT_Rhg1_ without epitope tags were co-expressed in *N. benthamiana* leaves under control of strong CaMV 35S promoters. GmRBOHC2 alone, AAT_Rhg1_ alone or GFP overexpression alone were expressed in the same leaf within the same biological replicate to serve as controls. 72 hrs after agroinoculation, leaves were detached, stained with nitroblue tetrazolium (NBT) for one half hour and then destained. NBT is a standard histochemical stain that detects superoxide (Beauchamp and Fridovich, 1971). Numerous NBT-positive spots were detected when coexpressing GmRBOHC2 and AAT_Rhg1_ or expressing AAT_Rhg1_ alone, while within the same leaves few or no NBT positive spots could be seen in the cells transiently expressing GmRBOHC2 alone, or GFP (Figure 9A). Quantification of staining areas confirmed a significant elevation of ROS production when expressing AAT_Rhg1_ alone. There was even more ROS production when GmRBOHC2 and AAT_Rhg1_ were co-expressed. Overexpression of GmRBOHC2 alone did not cause elevation of NBT staining beyond that observed for negative control GFP overexpression.

**Figure 9.**
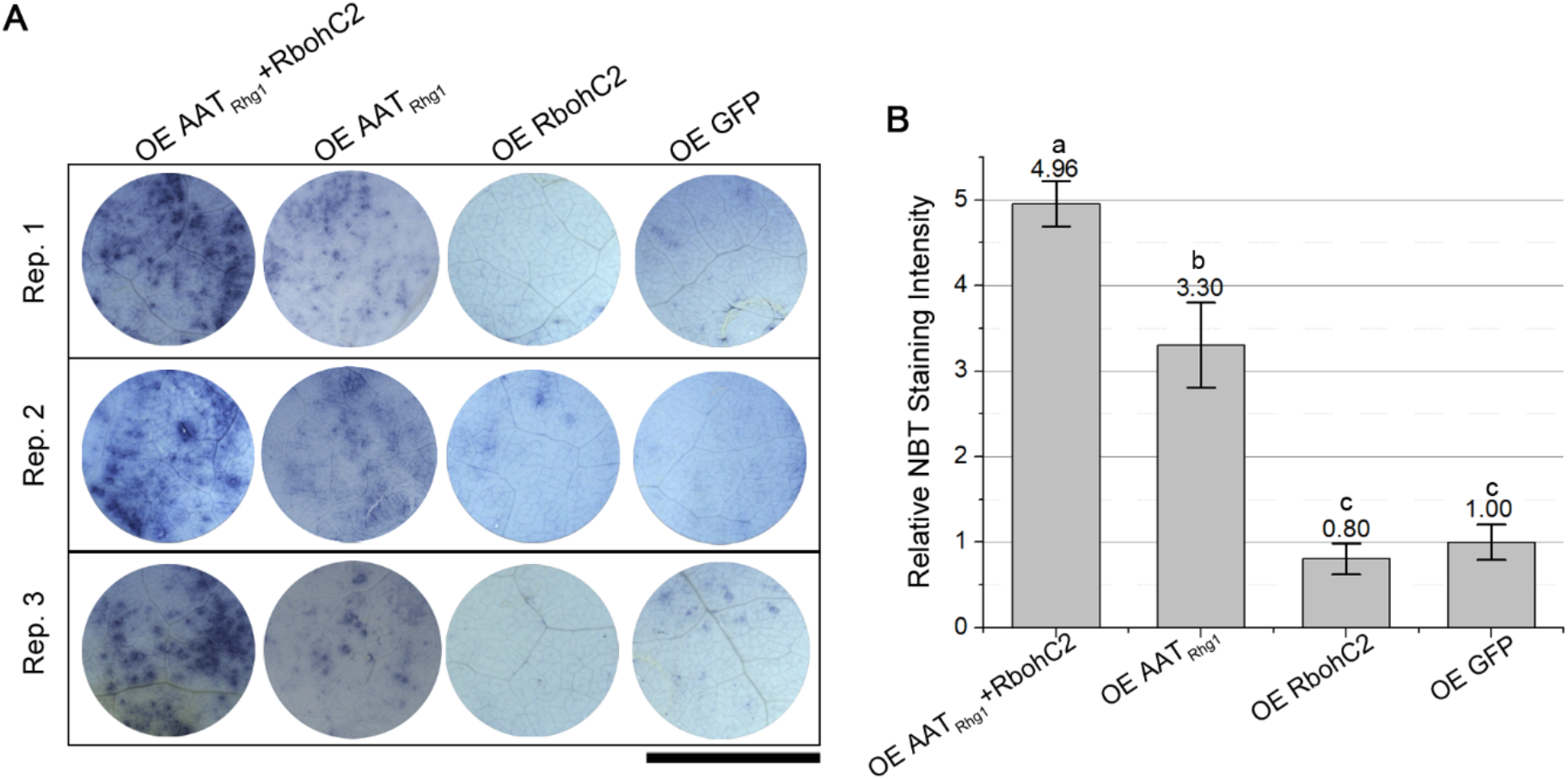
Simultaneous overexpression of GmRBOHC2 and AAT_Rhg1_ causes increased superoxide production in *N. benthamiana* leaves. **(A)** Representative images of *N. benthamiana* leaf regions stained with nitro blue tetrazolium (NBT) 72 hr. after agroinfiltration to overexpress (OE) (from left to right) GmRBOHC2 and AAT_Rhg1_ protein, AAT_Rhg1_ alone, GmRBOHC2 alone, or GFP control protein alone. Images from three biological replicates from separate dates are shown. Within each row, the images shown are all from the same leaf. Bar = 1 cm. **(B)** Quantification of NBT staining intensity of transformed leaf regions described in (A). Total area with NBT staining was measured using ImageJ and divided by total infiltration area, and normalized to the results for GFP alone control within the same replicate. n = 12 plants, mean ± SE shown, treatments with the same letter are not significantly different (ANOVA, p < 0.05).

## DISCUSSION

The *Rhg1* locus has for multiple decades been the primary means of control of the most economically damaging pathogen of soybean, soybean cyst nematode (Concibido et al., 2004; Niblack et al., 2006; Mitchum, 2016). *Rhg1* is a four-gene ~30kb block that exhibits copy number variation, and the PI 88788-type *rhg1-b* haplotype with 9-10 tandem *Rhg1* repeats is now present in most commercial soybean germplasm in the U.S. Contributions to SCN resistance have been shown for three of the four disparate genes in the *Rhg1* repeat block (Cook et al., 2012). Multiple mechanistic findings have been made regarding the resistance contributed by the α-SNAP_Rhg1_ proteins (Bayless et al., 2016; Lakhssassi et al., 2017; Liu et al., 2017; Bayless et al., 2018; Bayless et al., 2019; Lakhssassi et al., 2020). However, much less was known about how AAT_Rhg1_, or the Rhg1 protein with WI12-domains, contribute to resistance (Mitchum, 2016; Yan and Baidoo, 2018). One of the main findings of the present study is that AAT_Rhg1_ protein levels increase approximately ten-fold along the path of SCN root invasion. The hypothesis that AAT_Rhg1_ protein abundance differences are a determinant of resistance had previously been proposed (Cook et al., 2012; Cook et al., 2014), because unlike the α-SNAP_Rhg1_ proteins, there are no amino acid polymorphisms in AAT_Rhg1_ between resistant and susceptible varieties. Instead, *Glyma.18G022400* mRNA abundance in non-infected tissues had been shown to scale with *Rhg1* locus copy number, showing significant elevation in multi-copy *Rhg1* SCN-resistant genotypes (Cook et al., 2014; Wan et al., 2015). Moreover, before its function in SCN resistance was known, whole-genome expression profiling reported that *AAT_Rhg1_* mRNA abundance is elevated in SCN-infected root tissues (Kandoth et al., 2011; Matsye et al., 2011). It is interesting that by 7 dpi we observed similar levels of AAT_Rhg1_ protein abundance elevation, relative to neighboring cells, in both SCN-susceptible and SCN-resistant genotypes. However, at three days after SCN infection the abundance increase of AAT_Rhg1_ protein in cells along the SCN penetration path, relative to nearby non-infested cells, scaled with *Rhg1* locus copy number. *Rhg1* copy number has already been shown to positively correlate with SCN resistance efficacy, especially when isolated from contributions from other loci such as *Rhg4* (Cook et al., 2014; Yu et al., 2016; Patil et al., 2019). Our observations regarding AAT_Rhg1_ protein abundance increases are consistent with the documented differences in resistance efficacy between *Rhg1* haplotypes, but are striking with regard to the discovered site of AAT_Rhg1_ protein abundance increase.

The present study discovered that AAT_Rhg1_ protein specifically accumulates along the SCN root penetration path, relative to its abundance in the root cells adjacent to the penetration path or in any other observed root cells. In some of our study samples the SCN-penetrated root cells may have been dead or close to death at the time of fixation for microscopy, but they had obviously been stimulated to express elevated levels of AAT_Rhg1_ protein prior to that event. One of the reasons that this finding is of interest is because α-SNAP_Rhg1_, the protein product of the adjacent gene within the same *Rhg1* locus, was previously shown to accumulate more than ten-fold specifically within the syncytium cells that later serve for a few weeks as the biotrophic interface for cyst nematode feeding (Bayless et al., 2016; Bayless et al., 2019). We observed little or no AAT_Rhg1_ immunogold signal within syncytia for all three of the soybean varieties tested. This indicates that the resistance-contributing genes within the *Rhg1* locus not only encode distinctly different proteins, but those proteins also appear to act at spatially and temporally separate locations in the infection court. The multi-decade durability of *Rhg1*-encoded SCN resistance may have been due, in part, to the lower evolutionary potential of SCN relative to some microbial plant pathogens (McDonald and Linde, 2002). However, the present study provides experimental findings supporting the hypothesis that the durability of *Rhg1* is enhanced because it is a naturally occurring “resistance stack” encoding more than one mode of action.

The mechanism through which AAT_Rhg1_ activates defenses has remained unknown. None of the *Rhg1* genes encodes an NB-LRR, RLK or other protein type that commonly serves the role of pathogen detection and defense activation in plants (Dangl and Jones, 2001). Plants sense infection in one cell and then can activate defenses in systemic cells or nearby non-infected cells using a variety of mediators, including glutamate, ROS, Ca^2+^, salicylic acid, and N-hydroxypipecolic acid, to name just a few examples (Bernsdorff et al., 2016; Hartmann et al., 2018; Toyota et al., 2018; Wang et al., 2019). Local stimuli such as insect herbivory can activate glutamate as a longer-distance wound signal to rapidly initiate defense responses in undamaged parts (Toyota et al., 2018). ROS and electrical signaling, mediated by respiratory burst oxidase homolog (RBOH) proteins and glutamate receptor-like proteins, control distal activation of JA signaling during tomato responses to root knot nematodes (Wang et al., 2019). Those examples may be germane because AAT_Rhg1_, while not shown to be a glutamate transporter, was recently reported to increase tolerance to toxic levels of exogenously supplied glutamate, and it impacted glutamate abundance and transport (Guo et al., 2019). Our testing of a limited number of SCN infection-associated proteins revealed physical association of the AAT_Rhg1_ and GmRBOHC2 proteins in planta. We further observed that overexpression of AAT_Rhg1_ alone was sufficient to raise ROS levels above background, while co-expression of both proteins caused more elevation of ROS. We found that AAT_Rhg1_ abundance increases along the path of nematode invasion with greater 3 dpi increases observed in resistant haplotypes, and GmRBOHC2 mRNA abundance has been shown to increase in SCN-infected tissue (Wan et al., 2015; Liu et al., 2019). The present work hence establishes as a future priority a dissection of cause-effect relationships between AAT_Rhg1_/GmRBOHC2 interaction, ROS elevation and glutamate elevation in the activation of defense responses during cyst nematode infections.

Another intriguing feature of the present study was the extensive vesicle and macrovesicle production observed along the SCN penetration path in roots, and the association of AAT_Rhg1_ with vesicle and macrovesicle formation in *N. benthamiana* as well as along the SCN infection path in soybean. AAT_Rhg1_ expression induced vesicle/macrovesicle formation. In split-YFP experiments, GmRBOHC2 interaction with AAT_Rhg1_ was primarily observed on these vesicles. When considering the above and other findings about these AAT_Rhg1_-induced/AAT_Rhg1_-carrying vesicles, the vesicles do have things in common with the extracellular vesicles (EVs) recently found to play prominent roles during plant-microbe interactions (Rutter and Innes, 2017). First, they are not present in normal cells - their presence is elicited by pathogen infection. Second, they are in physically close contact with the penetration structure, for example, the nematode body (this study) or the haustoria structure (plant-fungal interaction). EVs that accumulate during microbial pathogen invasion have been shown to be defense cargo shuttle vectors, carrying various defense-related proteins, siRNAs and lipid signals that play roles in plant defense responses (Rybak and Robatzek, 2019). In Arabidopsis, defense related sRNAs could be shuttled into the necrotrophic fungus *Botrytis cinerea* via extracellular vesicles (Cai et al., 2018). The AAT_Rhg1_-associated vesicles that we observed may have similar roles during SCN infection. However, we have no evidence that the vesicles and macrovesicles are exported across the cell plasma membrane. They were observed within penetrated soybean root cells, or within *N. benthamiana* cells overexpressing AAT_Rhg1_, or in soybean root apoplastic fluid surrounding the nematode penetration path which includes intracellular remnants from recently deceased penetrated root cells. Significantly, in *N. benthamiana* FM4-64 experiments that monitored uptake of externally labeled plasma membrane, at least some of the AAT_Rhg1_-bearing vesicles were associated with endocytic rather than exocytic processes. In *N. benthamiana* overexpressing AAT_Rhg1_, co-localization of AAT_Rhg1_ with a plasma-membrane marker was observed on internally localized macrovesicles as well as the plasma membrane. Hence the observed AAT_Rhg1_-containing vesicles may be more reminiscent of the endocytic vesicles that become more abundant when RLKs such as FLS2 have been activated for signaling (Robatzek et al., 2006; Geldner and Robatzek, 2008; Beck et al., 2012).

ROS signaling plays important roles in plant responses to root knot nematodes (Zhou et al., 2018; Wang et al., 2019; Chen et al., 2020). In tomato, an oxidative burst had occurred by 12 hpi in both resistant and susceptible lines during root knot nematode infection. In electron microscopy, cerium chloride staining of H_2_O_2_ was mainly observed within cell walls near cells that underwent HR cell death caused by RKN infection (Melillo et al., 2006). ROS accumulation during beet cyst nematode infection was also reported in Arabidopsis (Waetzig et al., 1999; Siddique et al., 2014), and in both resistant and susceptible soybean roots upon SCN infection (Chen et al., 2020). However, detailed studies of the mechanisms that lead to ROS generation in soybean-SCN interactions are not available. We speculate, as one possibility, that the SCN-induced macrovesicles may provide a longer-lived membrane site for the ROS generation machinery. Within cells damaged by nematode penetration, these types of vesicles could serve as a briefly enduring cellular compartment where those defense responses can continue to function. A similar concept has been presented by Klink and colleagues, who proposed (Pant et al. 2015) that a transiently protected living plant cell could secrete materials in the vicinity of the nematode to disarm it, prior to that plant cell succumbing to it targeted demise. We observed that GmRBOHC2 physically interacts with AAT_Rhg1_ within vesicles. Upon SCN infection, the accumulation of AAT_Rhg1_ could recruit upregulated GmRBOHC2 onto those vesicles through their interaction. Alternatively, AAT_Rhg1_ may activate RBOHC2 and other respiratory burst oxidase homologs at the cell membrane, and then end up on vesicles simply as a recycling mechanism. The elevated coexpression of AAT_Rhg1_ and GmRBHOC2 did enhance ROS production, which may be a key early defense against SCN infection that directly weakens the nematode and/or signals to neighboring cells to potentiate defenses. It remains possible that defense-related functions other than ROS generation are also mediated by the observed AAT_Rhg1_-containing vesicles.

Taken together, we report distinct tissue and subcellular sites of elevated abundance of the putative amino acid transporter AAT_Rhg1_, along the path of SCN infection. AAT_Rhg1_ expression is associated with accumulation of vesicles and macrovesicles, and with activities that elevate ROS production, revealing mechanisms of the successful *Rhg1*-mediated SCN resistance that might be applied to other plant-nematode interactions.

## METHODS

### Nematode inoculum

SCN eggs of Hg 0 populations were obtained from Alison Colgrove at the University of Illinois Plant Clinic. Eggs were incubated in hatching buffer (3 mM ZnCl_2_) for 5 days at room temperature. Infectious J2 SCN were obtained and surface-disinfested with sterilization buffer (0.1g/L HgCl_2_ and 0.01% Sodium Azide) for three minutes. After rinsing twice in water, J2 SCN were resuspended in 0.05% sterile agarose water for root inoculation.

### Plasmid constructs

For transient overexpression vectors, the soybean AAT_Rhg1_ and GmRBOHC2 (*Glyma.06G162300.1*) ORFs were PCR-amplified from Williams82 cDNA generated by the iScript cDNA Synthesis Kit(Bio-Rad) and KAPA HiFi polymerase (Kapa Biosystems). Transient overexpression of soybean AAT_Rhg1_ and GmRBOHC2 was performed by assembling each respective ORF with the double CaMV 35S promoter with TMV omega enhancer (pICH51288) and nopaline synthase (NOS) terminator into the binary vector pAGM4673 (MoClo Tool Kit) using the Golden Gate cloning method (Weber et al., 2011).

For BiFC vectors, nYFP or cYFP fusion expression constructs driven by CaMV 35S promoter were prepared similarly as (Zhao et al., 2013). ORFs encoding AAT_Rhg1_ or α-SNAP_Rhg1_WT with stop codon, or GmRBOHC2 or NSF without stop codon were flanked by specific LIC1 adaptor at 5’(5’-C gAC gAC AAg ACC gTg ACC-3’) and LIC2 adaptor at 3’(5’-gA ggA gAA gAg CCg Tcg-3’) by overhang PCR amplification and purified by gel-extraction using QIAquick Gel Extraction Kit (Qiagen). Then, a Ligase Independent Cloning (LIC) method was performed to fuse N-terminal cYFP with AAT_Rhg1_ or α-SNAP_Rhg1WT_, and to fuse C-terminal nYFP with NSF or GmRBOHC2, as described (Xu et al., 2010).

For co-IP vectors, a construct encoding N-terminal GFP translationally fused to full length AAT_Rhg1_ (using an AAT_Rhg1_ cDNA with stop codon) was cloned into a binary expression construct pJG045, driven by a CaMV 35S promoter, using LIC (ligation-independent cloning) methods (Du et al., 2013). Similarly, a construct encoding N-terminal GmRBOHC2 without stop codon fused to C-terminal 6X MYC tag was cloned into pJG045 using LIC methods.

### *N. benthamiana* experiments

*N. benthamiana* plants with 2-3 fully expanded leaves were used for agroinfiltration with *Agrobacterium tumefaciens* strain GV3101(pMP90) as described in (Bayless et al., 2016). The subcellular compartment marker protein constructs expressed first 49 AA of GmMan1 (soybean α-1,2-mannosidase) as Golgi marker, chimeric signal peptide of AtWAK2 at the N-termnus of the RFP and the ER retention signal HDEL at the C-terminus as the ER marker, full length AtPIP2A as PM marker, and first 79 AA of small subunit of tobacco rubisco as plastid marker (Nelson et al., 2007)

### Antibody Production

Affinity-purified polyclonal antibodies, raised in rabbit against the synthetic peptide “SKGTPP” matching residues 15-20 near the N-terminus of GmAAT_Rhg1_, were produced by New England Peptide. Antibody specificity was validated using immunoblots with root lysates of ten-copy *Rhg1* Fayette) compared to single-copy *Rhg1* Williams 82 roots, and to Williams 82 roots expressing an RNAi gene silencing cassette targeting endogenous AAT_Rhg1_ (Figure S1).

### Immunoblots with anti-AAT_Rhg1_

Soybean root samples were frozen in liquid nitrogen and extracted in buffer containing 50 mM Tris·HCl (pH 7.5), 150 mM NaCl, 5 mM EDTA, 0.2% Triton X-100, 10% (vol/vol) glycerol, and protease inhibitor mixture (Sigma, P9599). Protein was extracted by homogenization in a PowerLyzer 24 (MO BIO) at 2000 rpm for three cycles with 15 s interval. Each sample was quantified by Bradford assays to achieve equal loading of total protein on SDS/PAGE gels. Blots were incubated with anti-AAT_Rhg1_ antibody in 5% (wt/vol) nonfat dry milk TBS-T (50 mM Tris, 150 mM NaCl, 0.05% Tween 20) at 1:1,000 overnight at 4°C. After four washes with TBS-T, secondary horseradish peroxidase-conjugated goat anti-rabbit was added at 1:10,000 and incubated for 1h at room temperature with mild agitation on a horizontal shaker. Blots were then washed with TBS-T four times, followed by chemiluminescence detection with SuperSignal West Pico or Dura chemiluminescent substrate (Thermo Scientific). Blots were imaged using a ChemiDoc MP chemiluminescent imager (Bio-Rad).

### Electron Microscopy and Immunolabeling

Immunolabeling were performed similarly to (Bayless et al., 2019). Segments from roots (Fayette, Forrest and Williams 82) previously inoculated with ~200 J2 SCN (Hg 0) per root were hand-sectioned with a razor at the indicated dpi. Root sections about 2 mm long were vacuum infiltrated in fixation buffer (0.1% glutaraldehyde and 4% (vol/vol) paraformaldehyde in 0.1M sodium phosphate buffer (PB) pH 7.4) and incubated overnight. After dehydration in 50%, 70%, 90%, 95% and 100% ethanol series, samples were embedded in LR White. Ultrathin sections (~90-nm) were taken longitudinally with an ultramicrotome (UC-6; Leica). For the immunogold labeling, samples were mounted on nickel slot grids. Grids were first activated on drops of 50 mM glycine/PBS for 15 min, and then blocked in drops of blocking solutions for goat gold conjugates (Aurion) for 30 min and then equilibrated in 0.1% BSA-C/PBS (incubation buffer). Next, grids were incubated with the anti-AAT_Rhg1_ antibodies diluted 1:1000 (in incubation buffer) overnight at 4 °C. After washing five times in incubation buffer, grids were incubated for 2 h with goat anti-rabbit antibody conjugated to 15-nm gold (Aurion) diluted 1:50 in incubation buffer. After six washes in incubation buffer and two washes in PBS, grids were fixed using 2.0% (vol/vol) glutaraldehyde in 0.1 M phosphate buffer for 5 min. Finally, grids were further washed twice in 0.1 M phosphate buffer for five minutes each and then five 2-minute washes in water. Images were collected with a MegaView III digital camera on a Philips CM120 transmission electron microscope. Anti-AAT_Rhg1_ immunogold particles were counted for single 69 μm^2^ areas within the sampled cells (e.g., cells penetrated by a nematode) and in the identically-sized region that had the highest observable signal in directly adjacent cells with normal root cell morphology (large central vacuole).

### Immunofluorescent assay

4 dpi SCN-infested roots segments were fixed in 0.1% glutaraldehyde and 4% (vol/vol) paraformaldehyde in 0.1M sodium phosphate buffer (PB) (pH 7.4) overnight after vacuum infiltration for about 1 hour as described above. For immunofluorescence processing, the fixed roots segments were briefly rinsed with PBT buffer (1XPBS pH=7.4, 1% BSA (w/v) and 0.1% Triton-X100) and then blocked with PBT blocking solutions (PBS pH=7.4, 1% BSA (w/v) and 0.1% Triton-X100, plus 5 % goat serum (Sigma-Aldrich) overnight at 4 °C. The root segments were incubated with the primary antibody diluted 1:1000 in PBT blocking solution at 4 °C overnight. Next, the incubated roots were washed 5 times for 10 min. each with PBT buffer at 800 rpm on a shaker platform at room temperature. Roots were then incubated with 0.4 μg/ml secondary antibody Alexa Fluor 568 goat anti-rabbit IgG H&L (Abcam ab175471) in PBT at for 2 hrs at room temperature, covered with a foil to shield the 2nd antibody solutions from light. After again washing with PBT for 10 mins 5 times at room temperature, the samples could be imaged by confocal microscopy right away.

### Confocal Microscopy

Confocal imaging was performed using an inverted Carl Zeiss laser-scanning confocal microscope (ELYRA LSM 780) with a 20× or 40× water immersion objective. All *A. tumefaciens*-transformed leaves were excised using a paper punch and monitored at ~72 hr after infiltration. For green fluorescent protein detection, GFP or GFP tagged chimera protein was excited at the wavelength of 488 nm, and the emitted fluorescence was detected with a 493-594 nm emission filter. Chloroplast autofluorescence was excited at 405 nm and detected at 635 to 708 nm to determine the position of chloroplasts for reference. For FM4-64 imaging, 50 μm FM4-64 solution (Invitrogen/Molecular Probes; T13320) was inoculated additionally into transformed leaves 0.5 hr before observation under the confocal laser scanning microscope. FM4-64 stained leaf tissues were excited with an excitation laser of 514 nm, and the emission signals were collected at 592-651 nm. For immunofluorescence, the Alexa Fluor 568 immunodetected plant tissue was excited at the wavelength of 561 nm and detection wavelength was at the range of 568-640 nm. For BiFC assay, YFP recombinant signal was acquired using a 514 nm laser for excitation combined with a 519-620 nm range emission filter. Images were collected using a standardized scan area of 442.2 × 442.2 μm (for 20× objective used) or 212.55 ×212.55 μm (40× water immersion objective), with a 1024 × 1024 pixels size frame. The detector master gain setting was from 700 to 800 dependent on different fluorescent signal intensity, and 1.01 AU size of pinhole was used for all the *Nicotiana benthamiana* samples, and 2.36 AU was used for soybean root immunofluorescence assays. At least 36 images were assessed for each expression treatment across three independent experiments.

### H_2_DCFDA detection of ROS in SCN-infested soybean roots

Whole 2-week old soybean seedlings germinated in PlantCon containers (MP bio, Cat#2672202) were used for SCN inoculation or mock treatment. About 400 SCN/root were placed near the vicinity of each root tip by pipette at day 0. After three days, the 2 cm root segments with greatest SCN infestation were harvested. For mock treatments, similar regions of the root were excised. Detached roots were then incubated in 1X PBS (Phosphate Buffered Saline) buffer with 50 μM H_2_DCFDA (2’, 7’-dichlorodihydrofluorescein diacetate, Invitrogen, D399) for 30 min shaking at 200 rpm/min at room temperature (Allan and Fluhr, 1997; Shin et al., 2005; Chen et al., 2020). Roots were then washed twice with 1X PBS, 10 min each time, and imaged. A Zeiss LSM 780 confocal microscope (ELYRA) was used with a 10× objective. H_2_DCFDA was excited at 488 nm at 2% laser power and 493-598 nm emission was detected. At least 16 confocal fluorescent images from 8 different roots across two independent replicates per treatment were used for quantification. The area with H_2_DCFDA fluorescent was calculated using ImageJ software as the number of pixels with signal intensity above background, compared to the total imaged root area (with SCN bodies and space outside the root tissue excluded) in order to calculate the percent area of root cells with ROS signals.

### MV treatment

Methyl Viologen (MV) treatment on *N. benthamiana* leaves was conducted as described (Han et al., 2015). In brief, 20 μm MV solution was infiltrated into the transformed leaves at 64 hr post agroinfiltration and followed by 8 hr under the previous light conditions to induce internal ROS generation before the confocal analysis.

### Coimmunoprecipitation

For co-IP assays, 4-wk-old fully expanded *N. benthamiana* leaves were used for agroinoculation at OD 0.6 for total 60-hr expression. About 2 g *N. benthamiana* leaf tissues for each treatment was collected from four different plants as one biological replicate. After chilling by liquid nitrogen, the tissue was ground by hand using a pestle in a pre-chilled mortar. Then 4 ml of protein extraction buffer (50 mM Tris·HCl (pH 7.5), 150 mM NaCl, 5 mM EDTA, 0.2% Triton X-100, 10% [v/v] glycerol, 1/100 Sigma protease inhibitor cocktail) was added to the mortar and the sample was further homogenized by grinding. The lysates were transferred into tubes and spun down at 6000 G for 10 min at 4°C three times to remove insoluble debris. The resulting supernatant was incubated with prewashed GFP-Trap_A (ChromoTek) beads for 3 h at 4°C. The precipitations were washed four times with ice-cold immunoprecipitation buffer at 4°C and were analyzed by immunoblot using anti-Myc (Sigma), or anti-GFP (Cell Signaling Technology) antibodies. Secondary horseradish peroxidase-conjugated goat anti-rabbit IgG (Sigma) was used to detect the primary anti-AAT_Rhg1_ antibody derived from rabbit. Chemiluminescence detection was performed with SuperSignal Dura chemiluminescent substrate (Thermo Scientific) and developed by a ChemiDoc MP chemiluminescent imager (Bio-Rad).

### Wounding treatment

For wounding, each *N. benthamiana* leaves was compressed gently for 30 seconds using reverse-action tweezers. By the full release of the reverse-action tweezers, a consistent wounding force was provided across all the samples.

### NBT staining

Nitro blue tetrazolium (NBT) staining was performed as described (Han et al., 2015). In brief, *N. benthamiana* leaves were detached and vacuum-infiltrated with 10 mM NaN_3_ in 10 mM potassium phosphate buffer (8.6 mM K_2_HPO_4_ and 1.4 mM KH_2_PO_4_) pH 7.8, for 1 min. Then, the fully infiltrated leaves were transferred into 0.1% NBT (in 10 mM potassium phosphate buffer pH 7.8) and put on a platform shaker shaking at 150 rpm for 30 min at room temperature, under a foil cover to reduce light exposure. The stained leaves were then cleared by boiling in destaining buffer (acetic acid: glycerol: ethanol (1:1:3 [v/v/v]). Photographs were obtained by scanning with a flatbed scanner (EPSON, V500 PHOTO SCANNER) at 800 dpi and NBT stain measurements were obtained using ImageJ (https://imagej.nih.gov/ij/). The region for evaluation was matched to the outline of the full agroinfiltrated area, and pixel intensities were obtained for each individual image. Number of dark blue-stained pixels in an image was divided by total analyzed pixels (the total infiltration area) to calculate the percent area with NBT stain. 32 images taken from 12 independent leaves across three independent replicates were used for quantification.

## Supplemental Data

**Supplemental Figure 1.** Confirming the specificity of custom-generated AAT_Rhg1_ antibodies.

**Supplemental Figure 2.** Little or no immunogold signal in mock-inoculated samples, syncytium cells, or negative controls that omit primary antibody.

**Supplemental Figure 3.** Confirming by competitive binding control that the AAT_Rhg1_ antibody is specific in electron microscopy antigen detection.

**Supplemental Figure 4.** AAT_Rhg1_ signal is not generated by microneedle damage

**Supplemental Figure 5.** GFP-AAT_Rhg1_ partially co-localizes with ER and PM markers but not with Golgi or plastid markers in *N. benthamiana* cells.

## ACKNOWLEDGEMENTS

We thank Dr. Ray Collier for sharing Golden Gate cloning parts and advice, Dr. Yule Liu and Dr. Jinping Zhao for sharing the LIC cloning vectors, and Ryan Zapotocny and Derrick Grunwald for helpful comments on the manuscript. This work was funded by the United Soybean Board and the Wisconsin Soybean Marketing Board.

## AUTHOR CONTRIBUTIONS

S.H. designed the research, performed research, analyzed data and wrote the paper. J.M.S. designed the research, performed research, and analyzed data. Y.D. performed research and analyzed data. A.F.B. designed the research, analyzed data and wrote the paper.

